# Transmembrane proteins tetraspanin 4 and CD9 sense membrane curvature

**DOI:** 10.1101/2022.06.02.494291

**Authors:** Raviv Dharan, Shahar Goren, Sudheer Kumar Cheppali, Petr Shendrik, Guy Brand, Li Yu, Michael M. Kozlov, Raya Sorkin

## Abstract

Multiple membrane shaping and remodelling processes are associated with tetraspanin proteins by yet unknown mechanisms. Tetraspanins constitute a family of proteins with four transmembrane domains present in high copy numbers in every cell type. Prominent examples are tetraspanin4 and CD9 that are required for the fundamental cellular processes of migrasome formation and fertilization, respectively. These proteins are enriched in curved membrane structures, such as cellular retraction fibers and oocyte microvilli. The factors driving this enrichment are, however, unknown. Here we revealed that tetrasapnin4 and CD9 are curvature sensors with a preference for positive membrane curvature. To this end we used a biomimetic system emulating membranes of cell retraction fibers and oocyte microvilli by membrane tubes pulled out of giant plasma membrane vesicles with controllable membrane tension and curvature. We developed a simple thermodynamic model for the partitioning of curvature sensors between flat and tubular membranes, which allowed us to estimate the individual intrinsic curvatures of the two proteins. Overall, our findings illuminate the process of migrasome formation and oocyte microvilli shaping and provide insight into the role of tetraspanin proteins in membrane remodelling processes.

## Introduction

Tetraspanins (TSPANs) are small proteins with four transmembrane domains present in every cell type^1,2^. TSPANs regulate cell morphology, adhesion, motility, fusion and signaling in diverse organs^3^. The functionality of TSPANs is thought to be dependent on their ability to form a distinct class of membrane domains by association among themselves and with other integral proteins and adhesion molecules^1,4^. TSPAN domains are dynamic and can change their composition and organization between different cell lines and cell states^5^.

Prominent members of the TSPAN family are TSPAN4 and CD9 whose crucial intracellular functions have been newly revealed. TSPAN4 has been shown to regulate formation of migrasomes, the recently discovered cell organelles playing essential roles in cell-cell signalling, lateral transfer of mRNA and proteins, transport of damaged mitochondria, and organ morphogenesis in vivo^6,7,8,9^. Structurally, migrasomes are few micron large spherical swellings formed on the retraction fibers, tens of microns long and about hundred nanometer thick protrusions of cell plasma membrane pulled out of the cell rear during migration^10^. Recent evidence suggests that retraction fibers are enriched with TSPANs and over-expression of 14 different TSPANs enhances migrasome formation^6^. Among them, tetraspanin 4 (TSPAN4) is one of the most effective. In the initial stage of migrasome biogenesis, migrasomes have relatively low TSPAN4 concentration. In the migrasome growing stage, however, TSPAN4 concentration in the migrasome membrane increases with time and was found to be essential for the mature migrasomes stabilization [Dharan *et. al*. 2022].

CD9 is involved in the viral infection cycle and localized to the growing tips of viral buds^11^. Furthermore, CD9 is imperative for the process of fertilization. Knockout of the CD9 gene resulted in abnormal distribution and shapes of microvilli in oocytes, which severely damaged the female fertility^12,13,14^.

While the biological roles of TSPAN4 and CD9 has been intensively addressed, the physico-chemical mechanisms underlying their functioning in cells remains unknown. Specifically, the physical forces underlying the observed affinity of these proteins to the strongly curved membranes of the retraction fibers (for TSPAN4) and microvilli (for CD9) have never been explored. At the same time, the existing structural data suggest that the background for this affinity may be related to sensing by these proteins of the membrane curvature. According to a recent crystallographic study^15^, CD9 has an asymmetric cone-like effective molecular shape and, hence, a positive intrinsic curvature^16,17,18^. As the transmembrane region is highly conserved for all TSPANs^15^, this whole family of proteins, including TSPAN4, might be characterized by considerable molecular intrinsic curvatures, which can provide a background for membrane curvature generation and sensing by these proteins in various processes^19,20,21,22^.

Here we revealed and quantitatively substantiated that TSPAN4 and CD9 are membrane curvature sensors. We experimentally demonstrated a large affinity of these proteins to membrane tubes having high positive curvature. We derived a thermodynamic model for protein enrichment in membrane tubes, which successfully described the strong partitioning of these proteins between nearly flat membranes of proteoliposomes and the strongly curved membrane tubules. Combining the experimental results and the modelling we estimated the effective intrinsic curvature of TSPAN4 and CD9.

## Results

We designed an artificial system of membrane tubules mimicking the cell retraction fibers and the shapes of oocytes-microvilli. TSPAN4 or CD9 with an N-terminal fluorescent GFP reporter were transiently expressed in HEK293T cells (Figure 1A, see methods for details). Giant plasma membrane vesicles (GPMVs) with TSPAN4 or CD9 in their membrane were immobilized on a glass surface within a custom-made chamber mounted on the stage of the correlated optical tweezers confocal fluorescence microscope (Figure 1A, B). An optically trapped polystyrene bead was pushed towards an immobilized GPMV and a membrane nanotube was pulled out of the vesicle (Fig 1C). The nanotube radius was significantly smaller and, hence, the curvature was considerably larger than those of the effectively planar GPMV. Measuring protein partitioning between the GPMV and the tubule allowed us to investigate the curvature sensing by TSPAN4 and CD9.

**Figure 1.**
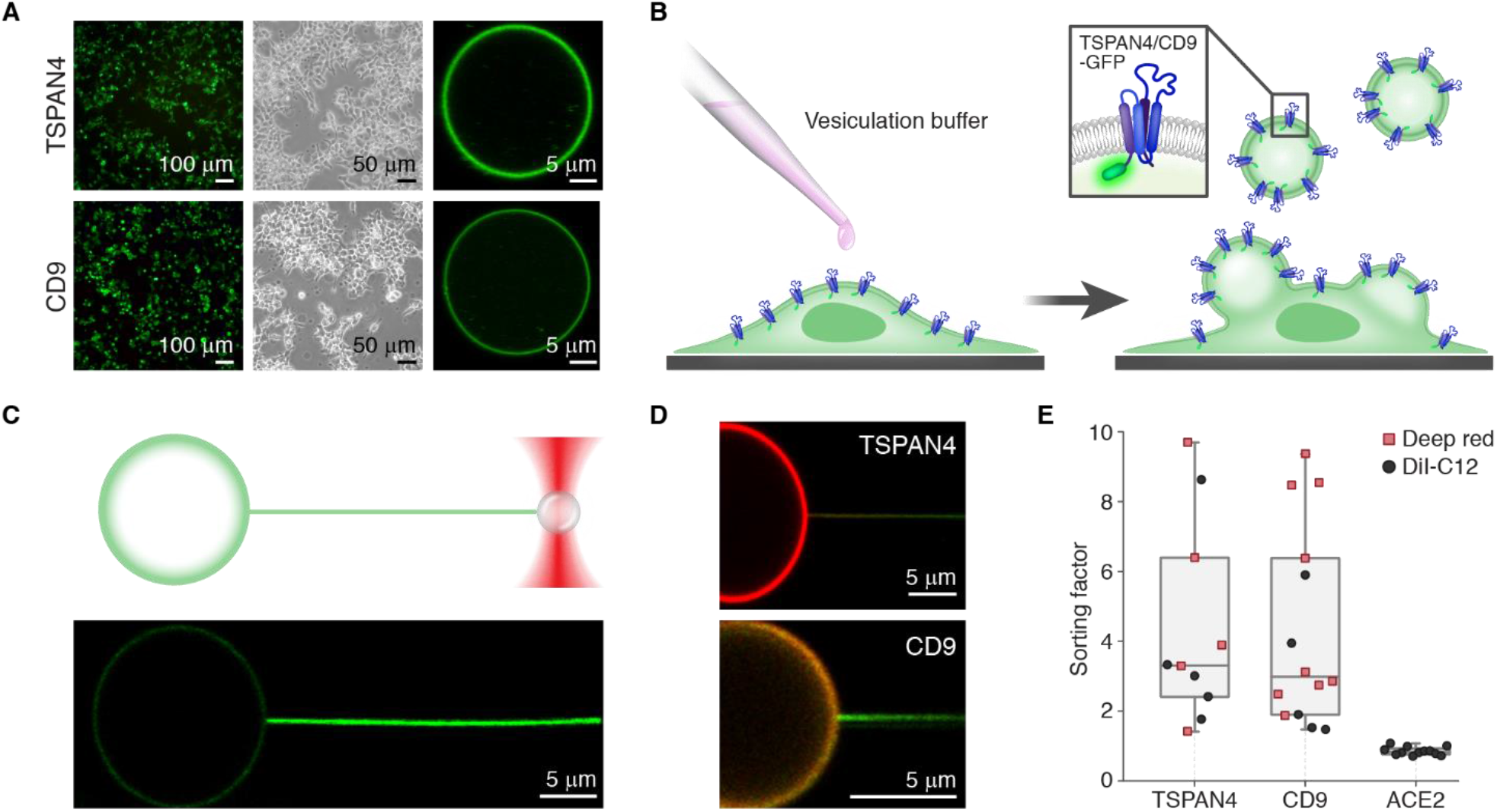
TSPAN4 and CD9 partitioning into curved membranes. (A) Microscopy images of HEK293T cells expressing TSPAN4\CD9-GFP (left images). After treatment with a vesiculation buffer, GPMVs can be seen floating in the sample or attached to the cells (middle images). The isolated GPMVs contained TSPAN4\CD9-GFP in their membrane (right images). (B) Illustration of the experimental procedure of GPMV formation. (C) Confocal microscopy image of a membrane tube pulled out of a GPMV containing TSPAN4-GFP. Top: schematic illustration of the tube pulling experiment, bottom: The tube was highly enriched with TSPAN4-GFP compared to the vesicle. (D) Confocal microscopy images of a tube pulled from a GPMVs containing TSPN4-GFP (top) or CD9-GFP (bottom) and labeled with the membrane dye DiI-C12 (GFP-green, DiI-C12-red). (E) Box plot comparing the sorting ratio of TSPAN4-GFP (n=10 vesicles), CD9-GFP (n=14 vesicles) and ACE2-GFP (control, n=12 vesicles). Black circles and red squares represent experiments conducted with DiI-C12 and Deep red, respectively.

After the tube was pulled, we monitored the time evolution of fluorescence of the GPMV and the tube (Fig. 1C). The fluorescence intensity of both TSPAN4-GFP and CD9-GFP at the membrane tube increased significantly with time, indicating that the proteins migrated to regions of higher membrane curvature (Figure 1C, S1A). Hence, membrane curvature can induce TSPAN4 and CD9 redistribution from the flat membrane of GPMV to the curved tubular membrane. In order to quantify this effect, we labelled the membranes by a plasma membrane dye (DiI-C12 or Deep red) by incubating the cells with the dye before generating GPMVs (Figure 1D, S1B). Protein enrichment could be measured by comparing its fluorescence intensity to that of the membrane dye^20,23^. If the affinity of TSPAN4-GFP\CD9-GFP to the tube is higher than to the flat membrane, the tube will appear green compared to the vesicle. On the other hand, if the affinity to the tube is the same or lower than to the flat membrane, the tube will have the same color as the vesicle or will appear redder (DiI-C12 or Deep red).

To quantify the protein redistribution to the curved membrane, we define the sorting ratio, *S*, as

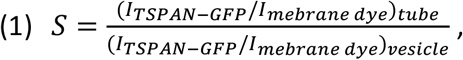

where *I* is the fluorescence intensity of the relevant component. According to this definition, a value of *S* greater than 1 indicates that the tube is enriched in TSPAN4\CD9. Our data showed that for both TSPAN4 and CD9, the sorting ratio was much higher than 1 (Figure 1E), demonstrating the striking sensitivity of these proteins to membrane curvature. A control experiment was performed with the transmembrane protein ACE2-GFP, which did not partition into the tube (Figure 1E, S1C). These high sorting ratios demonstrate that TSPAN4 and CD9 sense membrane curvature and partition into membranes with high positive curvature. The range of sorting ratios measured for TSAPN4 and CD9 was broad ranging between 1.5 and 10. A possible cause for this is a variability of the tube curvature, as in this assay the tube curvature was not controlled.

In order to confirm our hypothesis, we demonstrated that higher membrane curvatures induce higher TSPAN4\CD9 sorting. For this purpose, we integrated micropipette aspiration within dual-trap-tweezers confocal fluorescence microscope. The combination of micropipette aspiration with optical trapping allows one to control the membrane tension of the vesicle and thus the diameter of membrane tube, i.e., the membrane curvature of the tube, while measuring the pulling force of the tube (Figure 2A, B). We generated tensions in the vesicle membrane in the range 1.1 × 10^−5^ − 1.3 × 10^−4^ *N*/*m*. First, we pulled a tube upon a relatively low membrane tension, which resulted in a partitioning of TSPAN4\CD9 into the tube (Figure 2C). We then increased the tension in the vesicle, which led to higher pulling force (Figure S2) and concomitantly to higher tube curvature. This resulted in further protein enrichment in the tube (Figure 2C).

**Figure 2.**
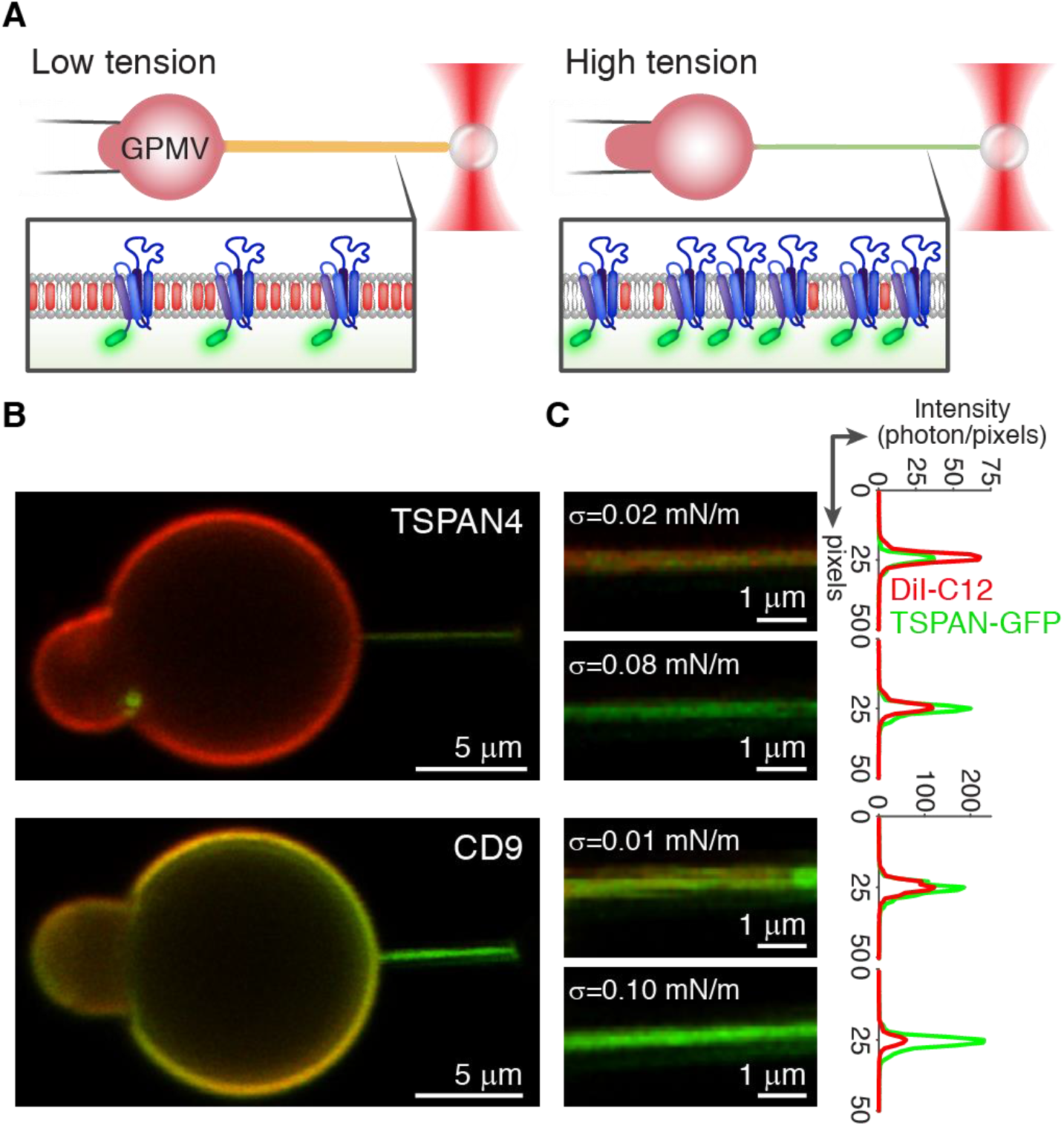
Higher membrane curvature increases TSPAN4 and CD9 sorting. (A) Schematic illustration of tube pulling with a controllable membrane tension setup. An optically trapped bead is used to pull a membrane tube from an aspirated GPMV by a micropipette. The pressure in the pipette determines the membrane tension and hence controls the tube radius. In parallel, the pulling force is measured while the GPMV and tube are scanned by confocal fluorescence microscopy and monitored in real time by bright field microscopy. (B) Confocal microscopy images of aspirated GPMVs containing TSPAN4-GFP or CD9-GFP (green) and labelled with DiI-C12 (red). (C) Confocal microscopy images of membrane tubes pulled from the GPMVs shown in B at relatively low and high membrane tension, as indicated. On the right the normalized intensity profiles of GFP and DiI-C12 of each image are presented.

To understand the physical factors determining the protein sorting in our system, we developed a thermodynamic model (see Supplementary Information). We assumed the membrane monolayers to be composed of lipid matrix with vanishing spontaneous curvature and to embed inclusions characterized by a positive molecular intrinsic curvature. We considered the system to consist of a flat membrane subjected to a constant tension and a tube (membrane tether) pulled out of the flat membrane by application of a local force. The model accounted for interplay between the bending energy of the tube membrane, the energy of the tension, and the entropy of the protein distribution in the membrane plane. Minimization of the system free energy enabled us to predict the dependence of the tube equilibrium radius, pulling force, and the difference between the protein concentrations in the tubular and flat membrane regions on the protein intrinsic curvature and the tension (see Supplementary Information). The free energy landscapes and model predications for different values of the model parameters are shown in Figures S3 and S4.

We next set out to fit our experimental results to the model predictions. This was done by numerically solving the model equations for various values of the membrane bending rigidity *κ* and the protein intrinsic curvature ζ with a goal to find the values of these two parameters providing the best agreement between the predicted and the experimentally determined values of the pulling force and sorting ratio as functions of the tension for each GPMV. The assumed parameters were the protein initial surface fraction (TSPAN4: 0.057-0.636%, CD9: 0.033-0.292%) and the in-plane areas of a lipid molecule and an inclusion taken, respectively, as 0.7 *nm*^2 24^ and 9.6 *nm*^2^, the latter based on the crystal structure of CD9^15^.

In most cases, the model predictions were in a good agreement with the measured values of the force and the sorting ratio (Figure 3A, B), yet for some of the experiments there were substantial deviations of the predicted values from the measured ones (Figure S5, S6). The fitting resulted in the mean values of the molecular intrinsic curvature of 0.11 ± 0.03 *nm*^−1^ for TSPAN4 and 0.12 ± 0.03 *nm*^−1^ for CD9 (Table S1 and Table S2).

**Figure3.**
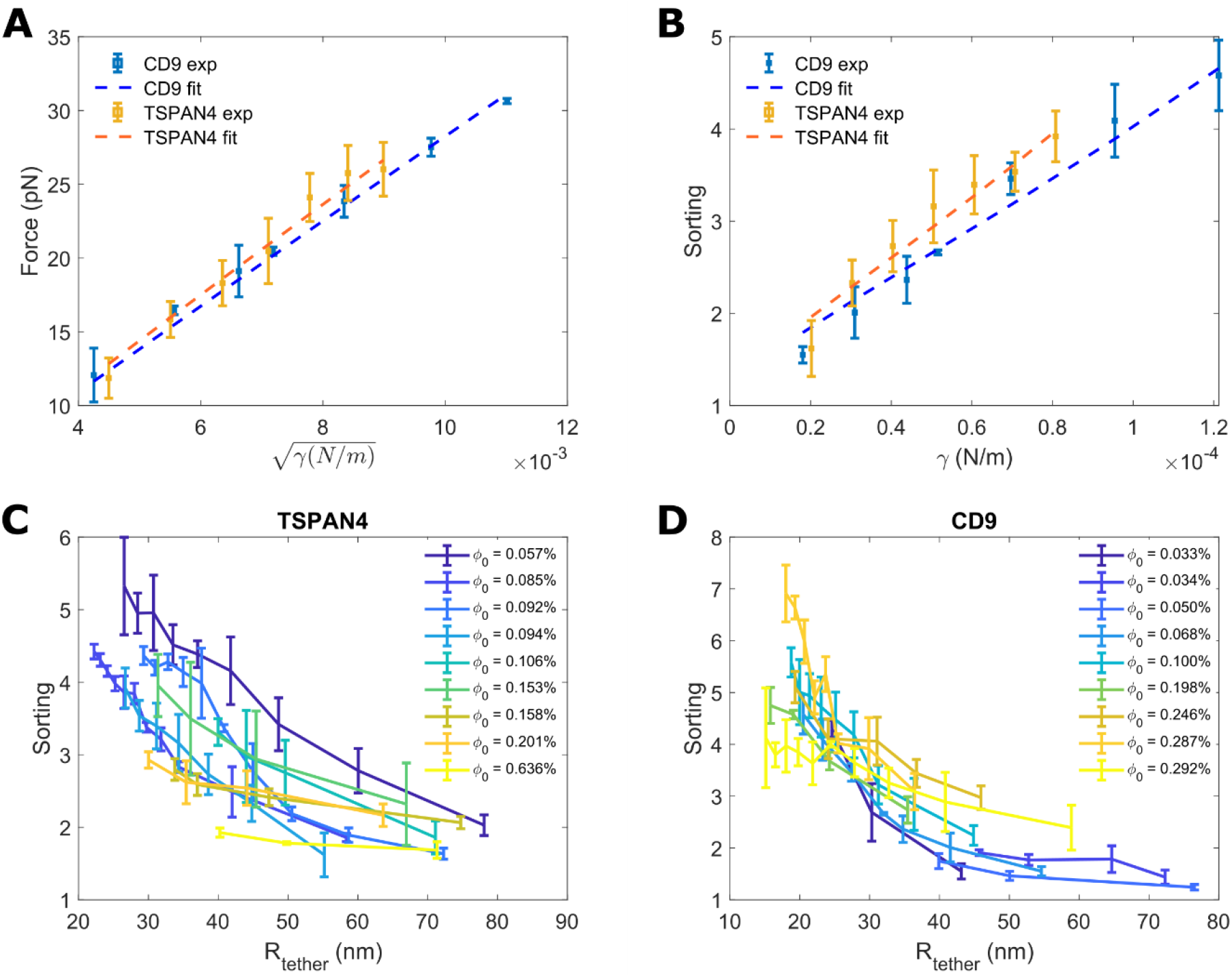
Free energy model for protein enrichment in membrane tube. (A, B) Plots for the pulling force as a function of square root of membrane tension (A) and sorting ratios as function of the membrane tension (B) of representative GPMVs containing TSPAN4-GFP or CD9-GFP. Squares are the mean force values; error bars are standard deviation. Dashed lines are fits of the numerical solutions for the free energy model with bending moduli *κ* = 32.935 ± 1.774 *k*_*B*_*T* and intrinsic curvature ζ = 0.120 ± 0.006 *nm*^−1^ for TSPAN4 or *κ* = 27.685 ± 0.988 *k*_*B*_*T* and ζ = 0.121 ± 0.007 *nm*^−1^ for CD9. (C, D) Sorting values of TSPAN4 (C) and CD9 (D) as function of membrane tube radius. Error bars are standard deviations.

The model predicted the sorting ratio to monotonically increase with increasing tension or, equivalently, decreasing tube radius. This trend for sorting as a function of the tether radius was observed experimentally for both TSPAN4 and CD9 (Fig.3B). Theoretically, the sorting ratio dependence on the protein concentration in the flat membrane playing a role of a proteins reservoir was predicted to be weak (Figure S4, H, J). This model prediction agreed with the experimental data for CD9-GFP (Figure 3D), whereas the sorting ratios measured for TSPAN4-GFP, seemed to be dependent on the reservoir protein concentration (Figure 3C).

The results described above correspond to the experiments in which the flat membrane tension was gradually increased from low to high values. For tubes, which did not rupture in the course of the tension growth, we investigated the evolution of the sorting ratio during a gradual decrease of the tension back to low values. On this backtrack, the sorting ratios were higher than those corresponding to the same values of the growing tension. Hence, sorting exhibited hysteresis (Figure S7), which was stronger for TSPAN4 than for CD9.

## Discussion

TSPAN proteins are ubiquitous in all cell types and are involved in diverse cellular phenomena^1,2^. The prominent examples are a critical role of TSPAN4 in the formation of migrasomes on retraction fibers^6^, and an essential involvement of CD9 in fertilization that occurs at the microvilli membrane of egg cells^12,13,14^. A common feature of the two processes is a preferential localization of TSPAN4 and CD9 to the strongly curved membranes of, respectively, the retraction fibers and micriovilli. The driving force leading to such localization remains unknown.

Here we discovered that both CD9 and TSPAN4 are membrane curvature sensors. Using a biomimetic system of membrane tubules with controlled radii pulled out of nearly flat membranes of GPMVs, we demonstrated a preferential partitioning of TSAPN4 and CD9 into membrane regions of high positive curvature. To quantitatively characterize the curvature sensitivity of these proteins we measured a correlation between the protein enrichment in the tubules as compared to the flat membranes in dependence on the tubular membrane curvature and quantified it by the sorting ratio. Based on these results we propose that the driving force for the retraction fibers and microvilli enrichment of TSPAN4 and CD9 is driven by the membrane curvature. To substantiate this proposal, we developed a simple thermodynamic model for protein partitioning between a flat membrane reservoir subject to a lateral tension and a tubular membrane pulled out of the reservoir by application of a local force. The model accounted only the interplay between the membrane curvature and the intrinsic curvature of the proteins and did not consider any protein-protein interactions.

While the model was able to recover the sorting ratio measured in most of the experiments, in a number of cases there were notable differences between the theoretical predictions and the experimental data. One main difference was observed for the experiments in which the flat membrane was subject to high tensions and, as a result, the tubular membrane reached large curvature values. In the large tension range, the measured sorting ratios, usually, reached a plateau as a function of the tension, whereas the model predicted a continuing increase in the sorting ratio. Another remarkable difference was that the measured sorting ratio of TSPAN4, but not CD9, exhibited a dependency on the reservoir protein concentration, whereas the model did not predict such dependency. Finally, while the model did not predict any hysteresis of the protein sorting ratio, for TSPAN4 the sorting ratios measured for a gradual increase on the tension were considerably lower than those observed during the tension decrease for the same tension values. For CD9 the observed hysteresis was very mild.

All of these differences between the observation and the theoretical predictions can likely be explained by interactions between TSPAN4 molecules, which have been previously described for TSPANs^1,2,3,4,5^. According to our results, these protein-protein lateral interactions are stronger for TSPAN4 than for CD9.

While our simple model does not account for the interactions between the TSPAN molecules, it nonetheless well explains the high sorting ratios of TSPAN4 and CD9 and their dependence on the tension in most on the tension range. Importantly, it allowed us to estimate the effective molecular intrinsic curvatures of the two proteins. The obtained high values of these intrinsic curvatures illuminate the strong influence of membrane shape on the localization and enrichment of TSPAN4 and CD9.

## Materials and Methods

### Cell culture, TSPAN4-GFP, CD9-GFP and ACE-GFP Expression Plasmids, cell transfection and giant plasma membrane vesicles (GPMVs) Isolation

HEK293T were cultured at 37°C and 5% CO_2_ in DMEM supplemented with 10% serum and 1% penicillin-streptomycin.

Complementary DNAs of tetraspanin 4, CD9 or ACE2 were cloned into pEGFP-N_1_. HEK293T cells were plated in 25 cm^2^ flask coated with poly-L-lysine (Sigma) to keep the cells attached during the blebbing process and to minimize cell debris in the solution. At 50% confluency, cells were transiently transfected with 5 μg DNA using Lipofectamine 2000 (Invitrogen) according to the manufacture’s protocols and then grown 24-36 hours for protein expression. GPMVs were produced according to a published protocol^25^. Briefly, following TSPAN4\CD9\ACE2-GFP expression, for the cells were stained with DiI-C12 (Invitrogen) or CellMask Deep red (Invitrogen) membrane dye, washed with GPMV buffer (10 mM HEPES, 150 mM NaCl, 2 mM CaCl2, pH 7.4) twice, and incubated with 1 mL of GPMV buffer containing 1.9 mM DTT (Sigma) and 27.6 mM formaldehyde (Sigma). Secreted GPMVs were then collected and isolated from the cells and immediately used for the optical trapping experiments.

### GUV preparation

Chloroform stock solutions of 99.9%%, 1,2-dioleoyl-sn-glycero-3-phosphocholine (DOPC, Avanti Polar Lipids) and 0.1% Oregon green 488 1,2-dihexadecanoyl-sn-glycero-3-phosphoethanolamine (Oregon green 488 DHPE, Invitrogen) were mixed at final lipid concentration of 0.25 mM. GUVs were grown on ITO slide (Nanion Technologies). 40 μL of 0.25 mM lipid solution in chloroform was gently spread with a needle on the slide, dried under argon gas and placed in a vacuum overnight.

GUVs were then grown by electroformation in 275 ml of 200 mM Sucrose solution, using Vesicle Prep Pro instrument (Nanion Technologies). The electroformation voltage was increased stepwise to 3 V (5 Hz) and performed at 37 °C for 2 hr.

### Tube pulling from immobilized GPMVs

The experiments were performed using a C-trap ® confocal fluorescence optical tweezers setup (LUMICKS) made of an inverted microscope based on a water-immersion objective (NA 1.2) together with a condenser top lens placed above the flow cell. The optical traps are generated by splitting a 10W 1064-nm laser into two orthogonally polarized, independently steerable optical traps. To steer the two traps, one coarse-positioning piezo stepper mirror and one accurate piezo mirror were used. Optical traps were used to capture polystyrene microbeads. The displacement of the trapped beads from the center of the trap was measured and converted into a force signal by back-focal plane interferometry of the condenser lens using two position-sensitive detectors. The samples were illuminated by a bright field 850-nm LED and imaged in transmission onto a metal-oxide semiconductor (CMOS) camera. **Confocal fluorescence microscopy:** The C-Trap uses a 3 color, fiber-coupled laser with wavelengths 488, 561 and 638 nm for fluorescence excitation. Scanning was done using a fast tip/tilt piezo mirror. For confocal detection, the emitted fluorescence was descanned, separated from the excitation by a dichroic mirror, and filtered using an emission filter (Blue: 500-550 nm, Green: 575-625 nm and Red: 650-750 nm). Photons were counted using fiber-coupled single-photon counting modules. The multimode fibers serve as pinholes providing background rejection. Experimental chamber: PDMS walls were placed on the bottom cover slip (Bar Naor) coated with poly-l-lysin (Sigma) and mounted onto an automated XY-stage. The GPMVs sample was added to the chamber and after about 15 minutes, a few drops of oil were put on the sample surface to prevent evaporation. In order to pull a membrane tube an optically trapped polystyrene bead (3.43 μm, Spherotech) was brought in contact with the GPMV for about a minute, and then moved away from the vesicle. For confocal imaging the 488, 532 and 638 nm lasers were used for GFF, DiI-C12 and Deep red excitation (respectively) with emission detected in three channels (Blue, Green, Red).

### Tube pulling from aspirated GPMVs

A micropipette aspiration setup including micromanipulator (Sensapex) holding a micropipette with diameter of 5 μm (Biological industries) connected to a Fluigent EZ-25 pump was integrated to our optical tweezers instrument. Before and after each experiment, the zero-suction pressure was found by aspirating a polystyrene bead into the pipette and reducing the suction pressure until the bead stopped moving. A membrane tube was pulled from aspirated GPMVs using beads trapped by the optical tweezers. First, a membrane tube was pulled at relatively low suction pressure (0.05-0.1 mbar, correspond to 1-2×10^−5^ N/m membrane tension), then we gradually increased the suction pressure (usually by 0.05-0.1 mbar) until we reached values in the range of 0.3-0.7 mbar (correspond to 5-13×10^−5^ N/m membrane tension), and then we gradually release back the pressure until reaching the zero pressure. At each suction pressure we monitored the force for at least 1 minute and scanned at least 3 confocal fluorescence scans (unless the tube was ruptured). For the confocal imaging the 488 nm and 532 nm lasers were used for GFP and DiI-C12 excitation with emission detected in two channels (Blue, Green).

### Data Analysis

Data acquisition was carried out using Bluelake, a commercial software from Lumicks. This software stores experimental data acquired during experiments with the C-trap in HDF5 files, which can be processed using Lumicks’ Pylake python package. Images of the confocal scans were reconstituted from photon count per pixel data in the HDF5 files using Pylake. All data analysis was performed with custom-written Python scripts. Fluorescence intensity profiles were obtained from the images by averaging the photon count of the relevant fluorescent channel in the region of interest.

### GPMV protein density

The GPMV protein-GFP density was determined based on published protocols^26,27^. Briefly, the fluorescence intensity of each GPMV was quantified according to the blue channel photon count of the confocal image in the region of interest. In separate experiments, giant unilamellar vesicles (GUVs) containing 0.1% Oregon green DHPE were imaged, which provided a reference signal of known fluorophore concentration in the membrane. 100 immobilized GUVs were scanned, at the same confocal setup, and the fluorescence intensity of the GUV was related to the fluorophore density using the equation 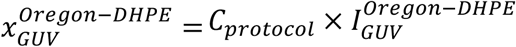 where *x* is the mole fraction of the fluorophore in the GUV, *C*_*protocol*_ depends on the confocal setup parameters and the fluorescence yield of the fluorophore, and *I* is the fluorescence intensity of the fluorophore. To compensate the fluorescence intensity yield between Oregon green and GFP, the intensity fluorescence of water solvated Oregon green and GFP were measured (Figure S8). The mole fraction of the protein in the GPMV was calculated by the equation:

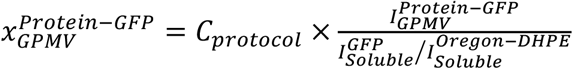

## Acknowledgements

RS acknowledges support by the ISRAEL SCIENCE FOUNDATION (grant No. 1289/20). SKC acknowledges support by the Ratner Center for Single Molecule Science. MMK was supported by Deutsche Forschungsgemeinschaft (DFG) through SFB 958 “Scaffolding of Membranes”, and Israel Science Foundation grant 3292/19, and holds Joseph Klafter Chair in Biophysics.

