## Supplementary Information for "Transmembrane proteins tetraspanin 4 and CD9 sense membrane curvature"

**Protein membrane tube enrichment model**

We consider a flat lipid membrane subject to lateral tension, $\gamma_0$. The membrane consists of lipid bilayer, which has bending rigidity, $\kappa$, and vanishing spontaneous curvature. The bilayer contains insertions modelling TSPAN molecules (or microdomains) and characterized by a molecular intrinsic curvature, $\zeta$. The insertions do not interact with each other and are free to distribute in the membrane plane by lateral diffusion. The molecular area of an insertion in the membrane plane, $a_p$, is larger than the molecular area of lipids, $a$, by a factor of $m$, so that $a_p = ma$. The area fraction occupied by the insertions in the flat membrane plane is $\phi_p$.

A point-like pulling force, $f$, is applied to the membrane and generates a cylindrical tube of length $L$. The insertions redistribute into the tube and their area fraction in the tube membrane is denoted by $\phi_t$. We assume the tube area to be much smaller than that of the flat membrane so that the latter plays for the former a role of a material reservoir with constant $\phi_p$.

Our goal is to determine the tubule radius, $R$, the pulling force, $f$, and the insertion area fraction in the tubule, $\phi_t$, as functions of the tension, $\gamma_0$, and the insertion area fraction, $\phi_p$, in the flat membrane reservoir. This will be done by minimizing the change of the system free energy (thermodynamic work) related to the tube formation.

We consider the thermodynamic work of the tube formation to have three contributions: the work, $F_\gamma$, of pulling the membrane area needed to form the tube, $A_t = 2\pi RL$, out of the flat reservoir against the tension, $\gamma_0$; the work, $F_B$, of bending the pulled membrane into the cylindrical shape of curvature, $J = 1/R$; the work, $F_S$, needed to change the insertion area fraction in the pulled membrane from the reservoir value, $\phi_p$, to that in the tube, $\phi_t$.

The overall energy change, $F_{tot} = F_\gamma + F_B + F_S$, will be minimized with respect to the tube radius, $R$, and the insertion area fraction in the tube, $\phi_t$.

Since the overall thermodynamic work must be independent of the pathway of the system transformations, we will consider the following sequence of stages: first, a spot of the initial membrane of area, $A_t$ , needed to form the tube, while remaining flat, undergoes a change of the insertion area fraction from $\phi_p$ to $\phi_t$; next, this spot is transformed into a cylindrical tube of cross-sectional radius, $R$, and length, $L$.

The free energy change corresponding to the first stage is $F_S$. Since no interaction between the insertions is considered by the model and the spot membrane remains flat in course of variation of the inclusion area fraction, the energy change, $F_S$, has a purely entropic character and can be calculated as

$$F_S = \frac{A_t}{ma}\int_{\phi_p}^{\phi_t}\left[\mu_t(\phi_t{}') - \mu_p\right]d\phi_t{}', \qquad \text{(S1)}$$

where

$$\mu_t(\phi_t') = k_B T\, ln\left(\frac{\phi_t{}'}{1-\phi_t{}'}\right) \text{ and } \mu_p(\phi_p) = k_B T\, ln\left(\frac{\phi_p}{1-\phi_p}\right) \qquad \text{(S2)}$$

are the entropic parts of the inclusion chemical potential in the spot membrane and the surrounding membrane, respectively.

The integration in (Eq.S1) accounting for (Eqs. S2) gives

$$F_S = k_B T\ \frac{A_t}{ma}\left[\phi_t\, ln\left(\frac{\phi_t(1-\phi_p)}{(1-\phi_t)\phi_p}\right) + ln\left(\frac{1-\phi_t}{1-\phi_p}\right)\right]. \qquad \text{(S3)}$$

The free energy change at the second stage of the flat spot transformation into a tubules includes the work of membrane bending, $F_B$, and of the membrane pulling out of the reservoir, $F_\gamma$.

To calculate the bending energy we use the Helfrich model of membrane elasticity [Helfrich 1973], according to which the variation of this energy, $dF_B$, resulting from a change of the membrane mean curvature, $J$, is given by

$$dF_B = A_t\kappa(J - J_s)dJ\ , \qquad \text{(S4)}$$

where $J_s$ is the spontaneous curvature of the membrane of the spot, which we assume to depend on the insertion area fraction according to Kozlov and Helfrcih[1],

$$J_s = \phi_t\zeta. \qquad \text{(S5)}$$

The bending energy change is then

$$F_B = \int_0^J A_t\kappa(J' - J_s)dJ' = A_t\kappa\left(\frac{1}{2R^2} - \frac{\zeta\phi_t}{R}\right), \qquad \text{(S6)}$$

where we used the relationship $J = 1/R$ for the cylindrical shape of the tube.

The energy of membrane tension corresponding to pulling out of the reservoir the area, $A_t$, consumed by the tube, is

$$F_\gamma = \gamma_0 A_t. \tag{S7}$$

Accounting for (Eqs.S3, S6, S7) and the relationship, $A_t = 2\,\pi R L$, the total energy change, $F_{tot}$, can be presented as

$$F_{tot} = 2\,\pi L R \left\{\kappa\left(\frac{1}{2R^2} - \frac{\zeta\phi_t}{R}\right) + \gamma_0 + \frac{k_B T}{ma}\left[\phi_t\, ln\left(\frac{\phi_t(1-\phi_p)}{(1-\phi_t)\phi_p}\right) + ln\left(\frac{1-\phi_t}{1-\phi_p}\right)\right]\right\}. \tag{S8}$$

Minimization of $F_{tot}$ (Eq.S8) with respect to $R$ and $\phi_t$results in the equilibrium values of these parameters, while the pulling force, $f$, is given by

$$f = 2\,\pi R \left\{\kappa\left(\frac{1}{2R^2} - \frac{\zeta\phi_t}{R}\right) + \gamma_0 + \frac{k_B T}{ma}\left[\phi_t\, ln\left(\frac{\phi_t(1-\phi_p)}{(1-\phi_t)\phi_p}\right) + ln\left(\frac{1-\phi_t}{1-\phi_p}\right)\right]\right\}, \tag{S9}$$

with $R$ and $\phi_t$ having the equilibrium values.

In the main text we use the results of numeric minimization of the energy (Eq. S8).

For the case of small differences between the inclusion area fractions in the tube and flat membranes, $\phi_t = \phi_p(1+\lambda)$ with $\lambda \ll 1$, the approximate results can be obtained analytically. After simple algebra one obtains

$$R = \sqrt{\frac{\kappa_{eff}}{2\gamma_0}}, \tag{S10}$$

$$f = \pi\left(2\sqrt{2\gamma_0\kappa_{eff}} - 2\kappa\zeta\phi_p\right), \tag{S11}$$

$$\phi_t = \phi_p\left[1 + \frac{ma}{k_B T}\kappa\zeta\left(1-\phi_p\right)\sqrt{\frac{\kappa_{eff}}{2\gamma_0}}\right]. \tag{S12}$$

where

$$\kappa_{eff} = \kappa(1 - \frac{ma}{k_B T}\kappa\zeta^2\phi_p\left(1-\phi_p\right)) \tag{S13}$$

is the effective bending modulus.

The consideration of insertions redistribution between a flat membrane and a cylindrical tube pulled out of it was presented in previous studies[2,3]. The results obtained here differ from those previous studies[2,3] since, according to our analysis and in contrast to that of previous studies[2,3], the contribution to the bending energy proportional to the spontaneous curvature square does not have to be included.

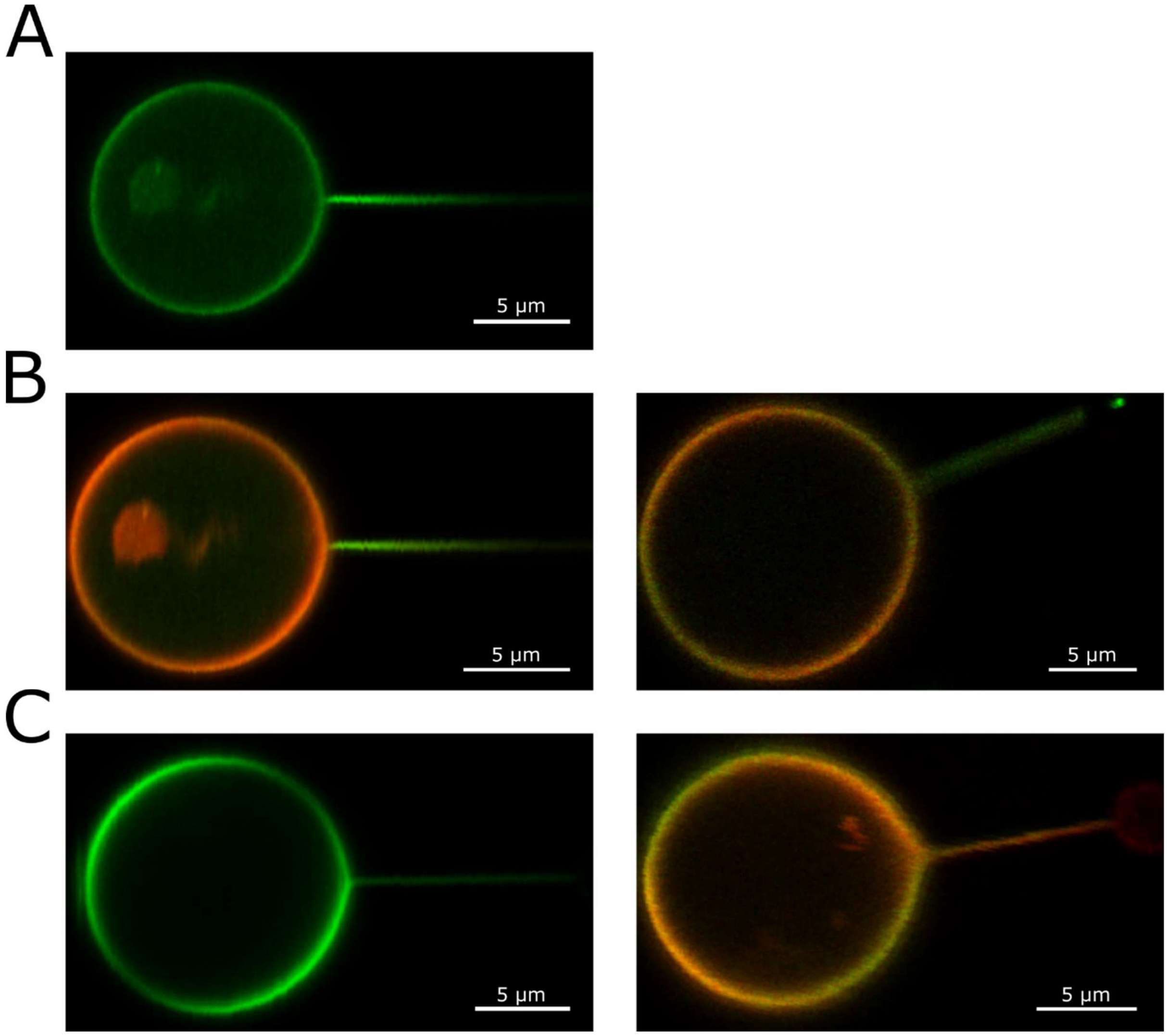


Figure S1. Membrane tubes pulled from giant plasma membrane vesicles (GPMVs). (A) Confocal microscopy image of a membrane tube pulled from GPMV containing CD9-GFP. The tube is highly enriched with CD9 compared to the vesicle. (B) Confocal microscopy images of a membrane tube pulled from GPMV containing CD9-GFP and Deep red (Left) or TSAPN4-GFP and Deep red (right, GFP-green and Deep red-red). (C) Confocal microscopy images of a membrane tube pulled from GPMV containing ACE2-GFP (left image) and ACE2-GFP and DiI-C12 (right image, GFP-green and DiI-C12-red). No tube enrichment was observed for ACE2-GFP experiments.

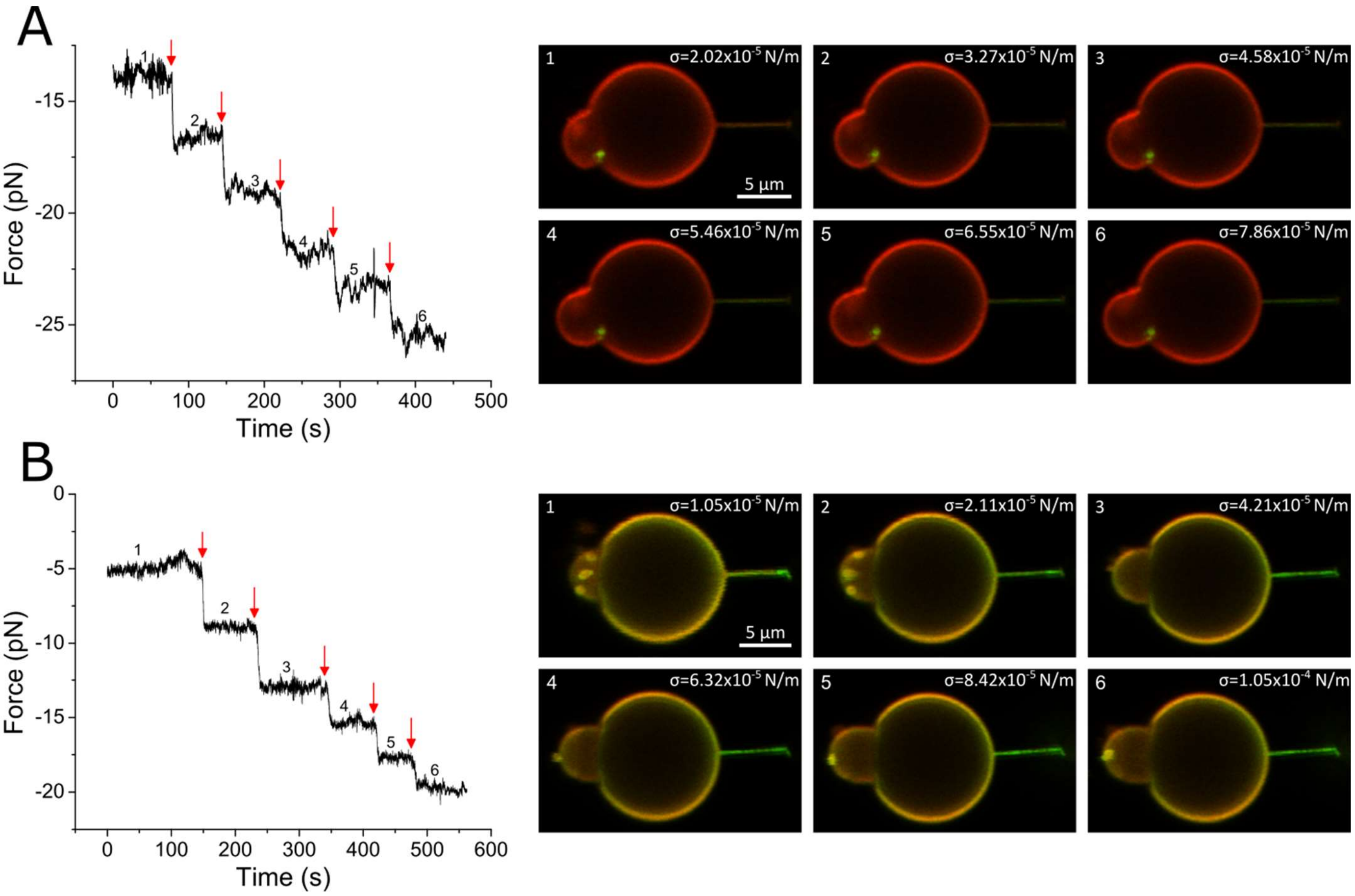


Figure S2. Tube pulling experiment. Force plots as function of the time of the force pulling a membrane tube from a GPMV containing TSPAN4-GFP (A) or CD9-GFP (B). Red arrows indicate tension increase and numbers corresponds to the images on the left.

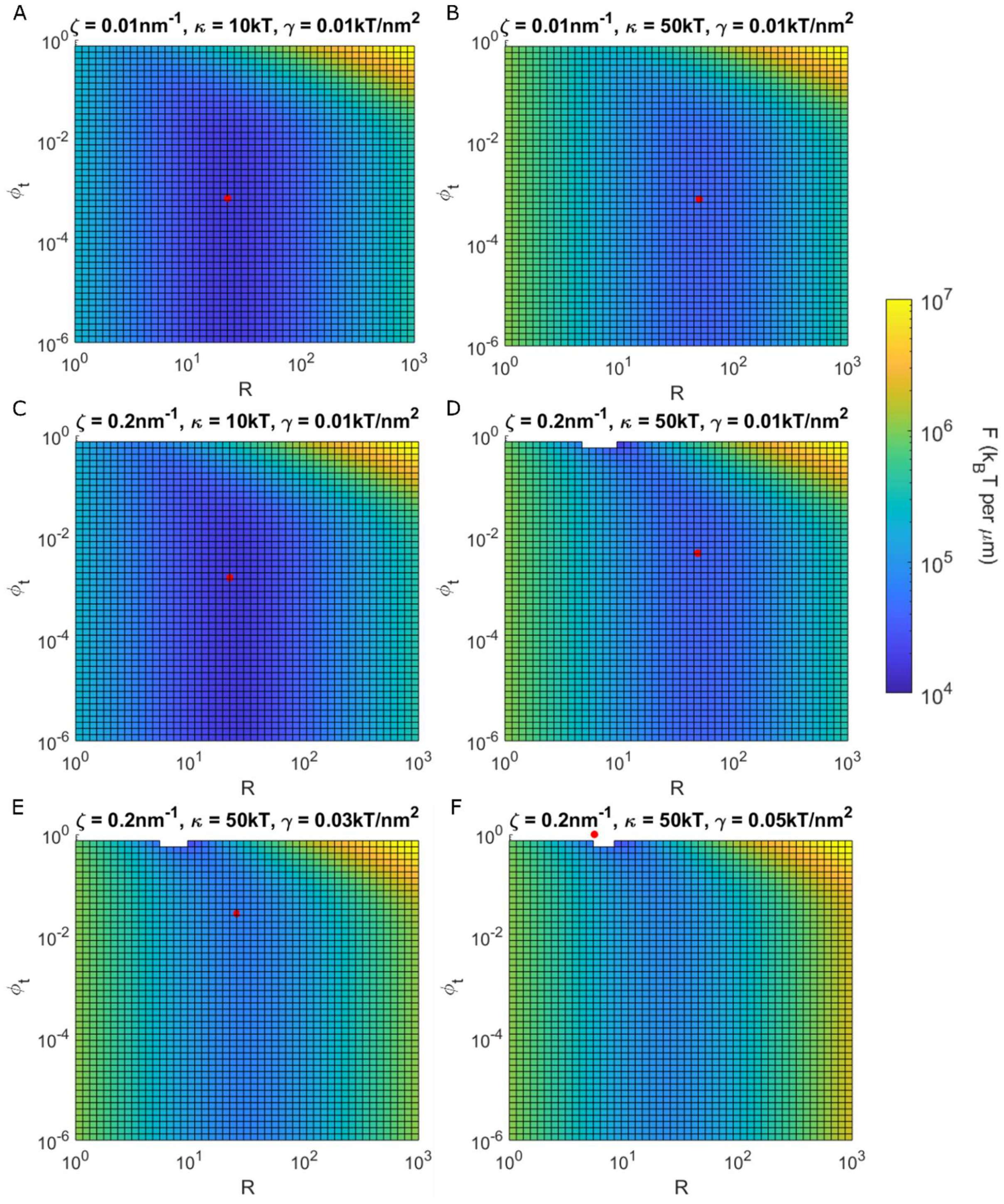


Figure S3. The model free energy $F$ as a function of membrane tube radius $R_{tet}$ and protein final area concentration $\phi_t$ are shown as color maps for different values of protein intrinsic curvature $\zeta$, membrane bending rigidity $\kappa$ and tension $\gamma$. The red circle corresponds to the free energy minimum. The white color areas shown in D-F correspond to negative energy values. In general, it can be seen that the free energy is more sensitive to $R_{te}$, which is mainly determined by as a competition between $\kappa$ and $\gamma$, with negligible influence of the protein on the tether geometry. On top of that, $\phi_t$ is determined by a competition between two free energy contributions: the mixing entropy, whose weight is proportional to the

$R_{tether}$, tends to keep $\phi_t$ low, while the protein-membrane curvature-mediated interaction, that becomes stronger with increased $\kappa$ and $\zeta$, pushes for high protein concentrations. In extreme cases (As seen in F), where both $\kappa$ and $\zeta$ are high, there is a critical value of $\gamma$, above which $\phi_t$ approaches 1 and $R_{tet}$ shrinks dramatically. This behavior is probably prevented in practice by some forces of short-range repulsion interaction between the proteins.

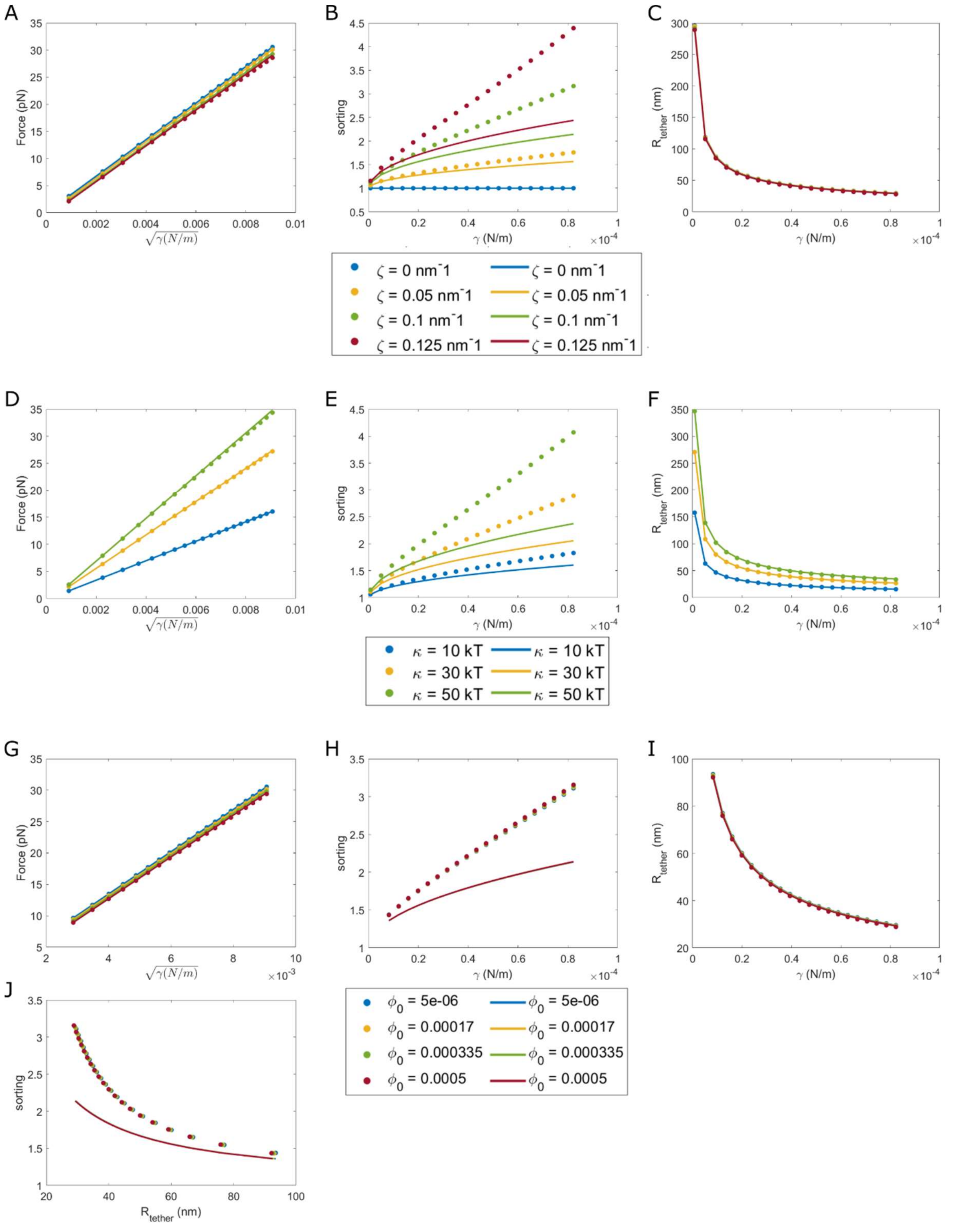


Figure S4. Model predictions are presented as plots of the pulling force, protein sorting and membrane tube radius predicted by the model as a function of membrane tension, for different values of protein intrinsic curvature, membrane bending rigidity and initial protein concentration. In all plots, circles correspond to numeric solutions obtained by minimization of the full free energy term, and solid lines correspond to analytical expressions in the limit of weak sorting. (A) Pulling force as function of the square root of membrane tension for different values of intrinsic curvature. It can be seen that the pulling force weakly depends on the intrinsic curvature, but for high values it can be slightly reduced, while maintaining the same slope. (B) Protein sorting vs. tension for different values of intrinsic curvature. The protein sorting dramatically increases with tension, and it is very sensitive to the intrinsic curvature. For small values of intrinsic curvature, the analytical predictions are in agreement with the numerical results, but for high sorting values they don't agree, as expected. (C) Tube radius as a function of tension for different values of intrinsic curvature. The radius is decreased with increasing tension, and is not sensitive to the protein intrinsic curvature. (D) Pulling force as function of the square root of membrane tension for different values of membrane bending rigidity. Membrane rigidity is the main parameter determining the slope of the force trend. (E) Sorting as function of the tension for different values of membrane bending rigidity. Higher bending rigidity leads to higher sorting values, due to the stronger protein-membrane curvature-mediated interactions. (F) Tube radius as a function of tension for different values of membrane bending rigidity. High bending rigidity increases the energetic cost of tube narrowing, and makes the tether somewhat wider. (G – J) The same trends as in A-C and D-E for different values of initial protein concentration. It can be seen that for values in the experimental range, the initial concentration does not change the results, besides slightly reducing the pulling force.

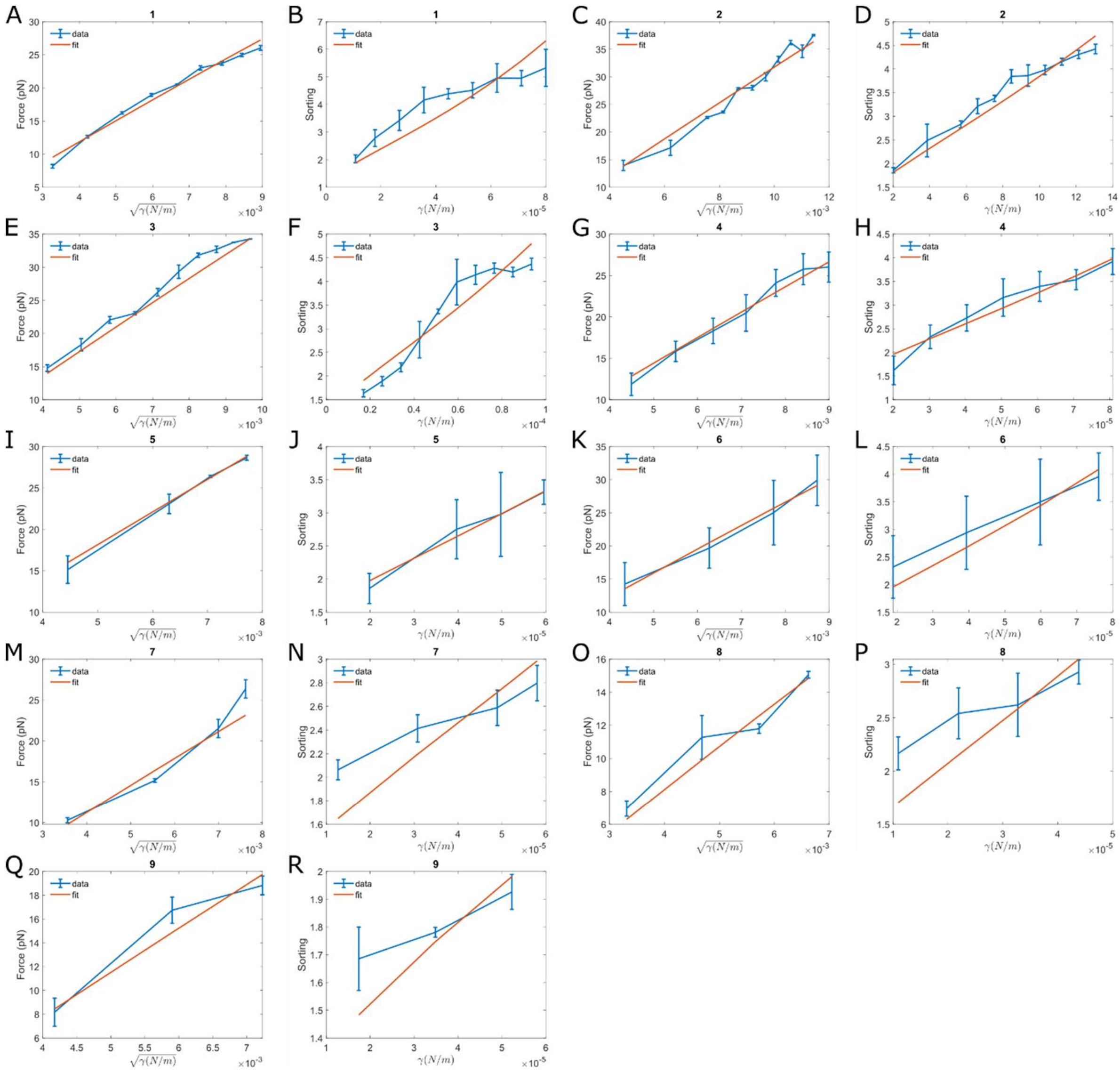


Figure S5. Numerical solutions for the minimization of the free energy fitted to the experimental measurements of membrane tube pulling from aspirated GPMV containing TSPAN4. Each pair of plots (A+B, C+D etc.) shows the model fit for force and sorting measurements. Numbers in the title of the figures are labels given to each vesicle for later identification.

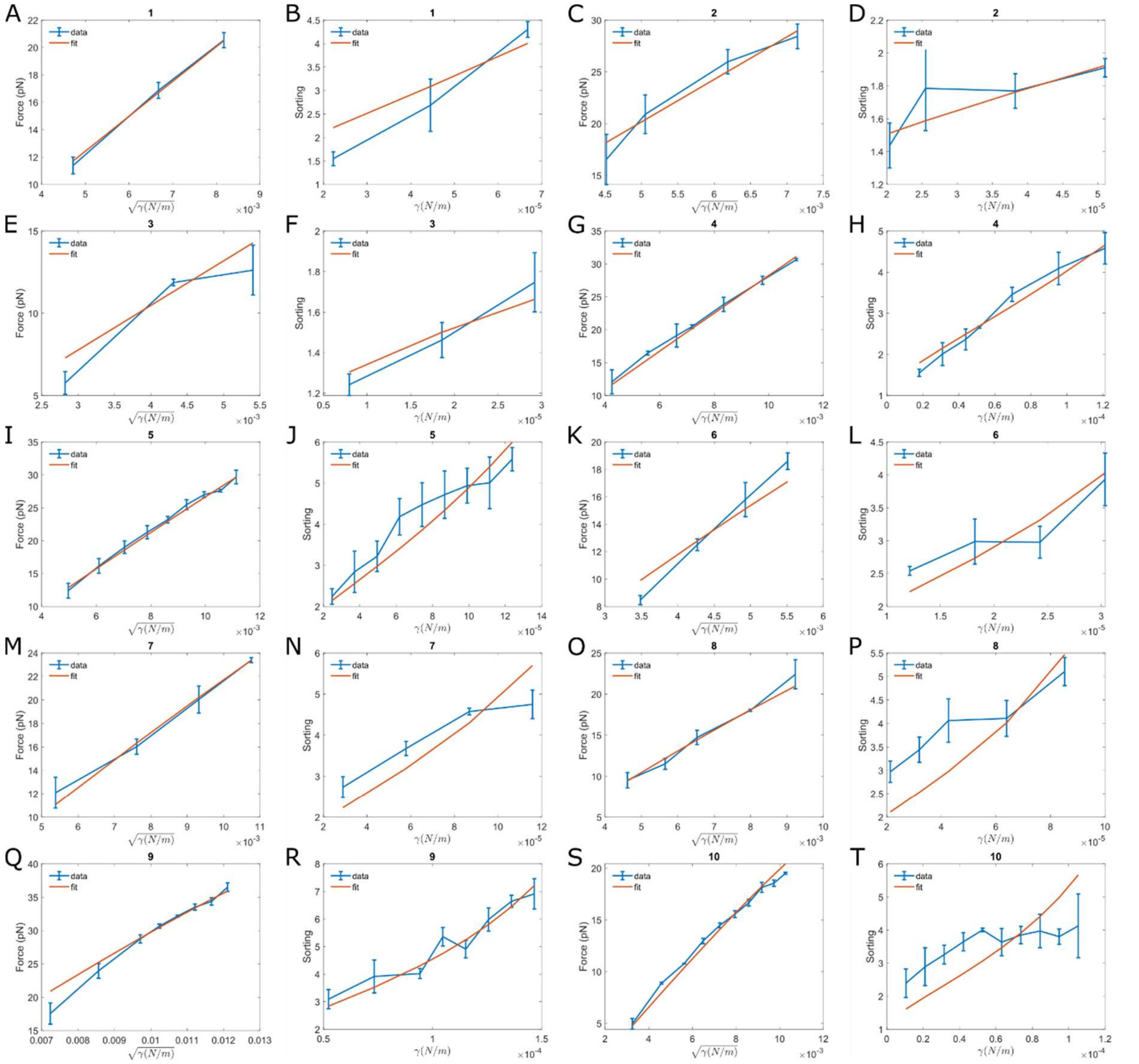


Figure S6. Numerical solutions for the minimization of the free energy fitted to the experimental measurements of membrane tube pulling from aspirated GPMV containing CD9. Each pair of plots (A+B, C+D etc.) shows the model fit for force and sorting measurements. Numbers in the title of the figures are labels given to each vesicle for later identification.

Table 1. Bending rigidity $\kappa$ and protein intrinsic curvature $\zeta$ used for fitting the model to the experimental data of TSPAN4

| Vesicle Number | κ (kT) | κ error | ζ (nm^-1) | ζ error |
|---|---|---|---|---|
| 1 | 34.769 | 1.829 | 0.150 | 0.009 |
| 2 | 35.773 | 1.787 | 0.103 | 0.007 |
| 3 | 47.693 | 2.046 | 0.104 | 0.008 |
| 4 | 32.935 | 1.774 | 0.120 | 0.006 |
| 5 | 54.575 | 1.93 | 0.0943 | 0.005 |
| 6 | 48.361 | 4.713 | 0.100 | 0.006 |
| 7 | 39.043 | 9.774 | 0.103 | 0.022 |
| 8 | 26.511 | 5.362 | 0.139 | 0.013 |
| 9 | 53.547 | 24.090 | 0.061 | 0.0073 |

Table 2. Bending rigidity $\kappa$ and protein intrinsic curvature $\zeta$ used for fitting the model to the experimental data of CD9

| Vesicle Number | κ (kT) | κ error | ζ (nm^-1) | ζ error |
|---|---|---|---|---|
| 1 | 21.110 | 3.119 | 0.170 | 0.029 |
| 2 | 52.519 | 4.156 | 0.060 | 0.014 |
| 3 | 23.040 | 6.214 | 0.093 | 0.094 |
| 4 | 27.685 | 0.988 | 0.121 | 0.007 |
| 5 | 27.142 | 0.876 | 0.134 | 0.004 |
| 6 | 59.601 | 16.164 | 0.123 | 0.012 |
| 7 | 21.822 | 2.244 | 0.142 | 0.007 |
| 8 | 29.095 | 3.433 | 0.129 | 0.006 |
| 9 | 46.837 | 1.811 | 0.091 | 0.001 |
| 10 | 22.519 | 2.594 | 0.137 | 0.005 |

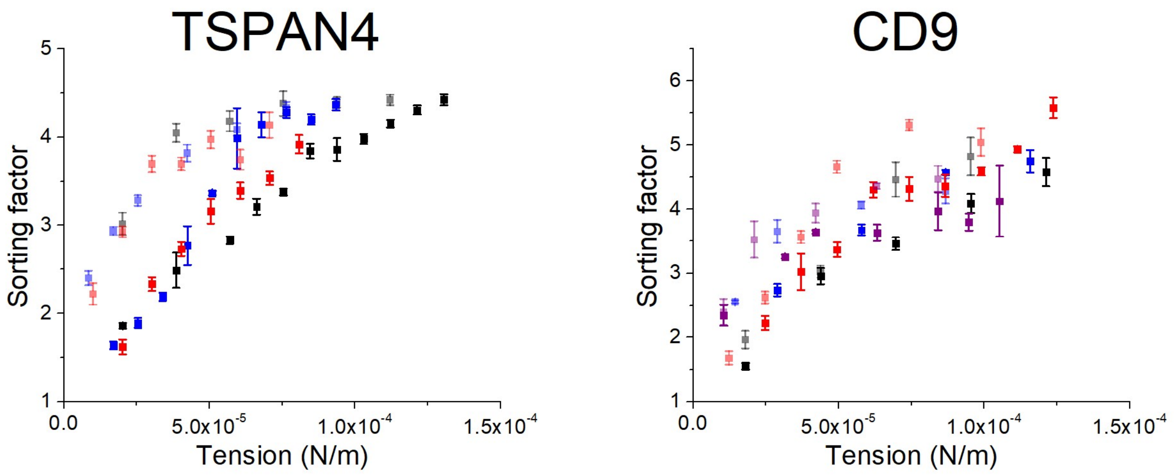


Figure S7. Hysteresis of TSPAN4 and CD9. Membrane tube was pulled from aspirated GPMV, which contained TSPAN4-GFP or CD9-GFP and the membrane dye DiI-C12, at relatively low tension. Next, by increasing the aspiration pressure the membrane tension was increased, which resulted in higher TSPAN4\CD9-GFP tube enrichment i.e., higher sorting factor values. When the aspiration pressure was drop back, the sorting values were higher for the same membrane tension values. Dark color squares represent tension increase path whereas light color squares represent tension decrease (n=3 vesicles for TSAPN4, n=4 vesicles for CD9). Error bars are SEM.

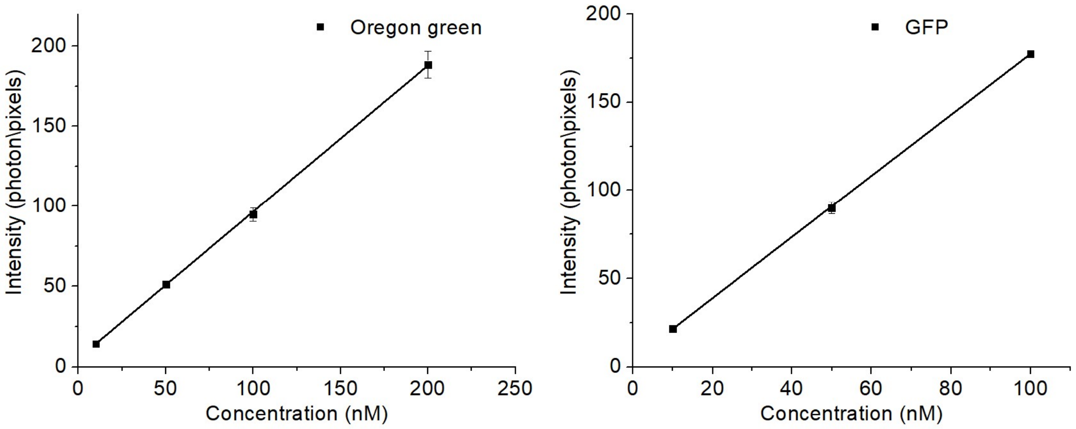


Figure S8. Normalized fluorescence intensity as function of the concentration of water-solvated Oregon green or GFP. Each point represents 10 scans, error bars are the sample standard deviation.
